# *Bdellovibrio* and Like Organisms are Predictors of Microbiome Diversity in distinct Host Groups

**DOI:** 10.1101/627455

**Authors:** Julia Johnke, Sebastian Fraune, Thomas Bosch, Ute Hentschel, Hinrich Schulenburg

## Abstract

Biodiversity is generally believed to be a main determinant of ecosystem functioning. This principle also applies to the microbiome and could consequently contribute to host health. According to ecological theory, communities are shaped by top predators whose direct and indirect interactions with community members cause stability and diversity. *Bdellovibrio* and like organisms (BALOs) are a neglected group of predatory bacteria that feed on Gram-negative bacteria and can thereby influence microbiome composition. We asked whether BALOs can predict biodiversity levels in microbiomes from distinct host groups and environments. We demonstrate that genetic signatures of BALOs are commonly found within the 16S rRNA reads from diverse host taxa. In many cases, their presence, abundance, and especially richness are positively correlated with overall microbiome diversity. Our findings suggest that BALOs can act as drivers of microbial alpha-diversity and should therefore be considered as candidates for the restoration of microbiomes and the prevention of dysbiosis.

---

Biodiversity is a key attribute of productive [1] and stable ecosystems [2]. This is likely due to the activity of highly productive keystone species [3], which are often more common in species-rich communities [1]. Nevertheless, productivity and stability appear to be mainly driven by diversity itself and not by individual taxa [4]. Species-rich communities exist for example in the human gut and oral microbiome and are usually assumed to consist of functionally redundant species that act as insurance in case of extinctions [5, 6]. Consequently, species-rich communities are more resilient (cf. [7]). To date, most studies on the effect of biodiversity on ecosystem functioning and specifically the effect of microbiome composition on host health have focused on a single trophic level. Yet, changes in the diversity of one trophic level can affect other trophic levels, either directly through consumer-resource interactions or indirectly when the decrease of one species leads to abundance changes of other species [8]. The presence of top predators has particularly strong effects because they can limit dominant species abundance and thereby free niches for rare taxa [9–11]. The impact of predators is likely distinct from environmental stressors, which may similarly free niches and subsequently increase microbiome diversity, as recently documented for the microbiome of *Daphnia* waterfleas after antibiotic exposure [12]. Yet, in this case, the effect on community composition is likely to be random, whereas predators usually target the dominant species.

*Bdellovibrio* and like organisms (BALOs) are obligate predators of Gram-negative bacteria in a wide range of habitats [13, 14]. BALOs were recently linked to a healthy human gut microbiome [15], and proposed as living antibiotics in medical treatment [16] and water remediation [17]. Additionally, a microcosm experiment showed that their predatory activity can exceed phage-induced mortality [18]. We here draw attention to this neglected group of predators and tested their association with microbial diversity as an indicator of a healthy microbiome across distinct animal host groups and environments.

We analyzed 16S rRNA data from randomly chosen, exemplary host taxa that are representative of distinct animal taxonomic groups, including early branching metazoans, ecdysozoa, selected vertebrates, and additionally home surfaces (Table S1 and Supplementary Methods). We only considered studies, if they included samples with and without BALOs, thereby allowing us to determine the consequences of BALO presence and absence in comparable groups. We determined BALO occurrence (although not necessarily activity) by identifying OTUs that showed 97% sequence identity to members of the BALO-containing taxonomic groups *Bdellovibrionales* (including the families *Bacteriovoracaceae* and *Bdellovibrionaceae*) and *Micavibrionales* (including *Micavibrionaceae*). From these data, we inferred relative BALO abundance and corresponding microbiome alpha- (i.e., Shannon-Wiener diversity, Simpson’s diversity, richness) and beta-diversities.

The presence of BALOs was associated with a significantly higher Simpson and Shannon diversity for the microbiomes of seven and five host species, respectively, as well as the home surfaces (Figure 1, Table S2). The main exceptions referred to two sponge species, *Carteriospongia foliascens* and *Ircinia variabilis*, which showed a significantly higher alpha-diversity in the absence of BALOs. This negative association was not observed for microbiome richness (Figure S1, Table S3). Our subsequent analysis of absolute OTU numbers revealed that microbiome richness is significantly associated with both BALO abundance (Figure 2a, Table S4) and BALO richness (Figure 2b, Table S4) in case of *H. vulgaris* and the sponges. A trend toward this association was additionally observed for *N. vectensis* and *D. melanogaster*. Interestingly, for both host systems, OTU richness was highest with medium BALO abundance, which possibly indicates that BALO richness rather than abundance influences microbiome richness.

**Fig. 1.**
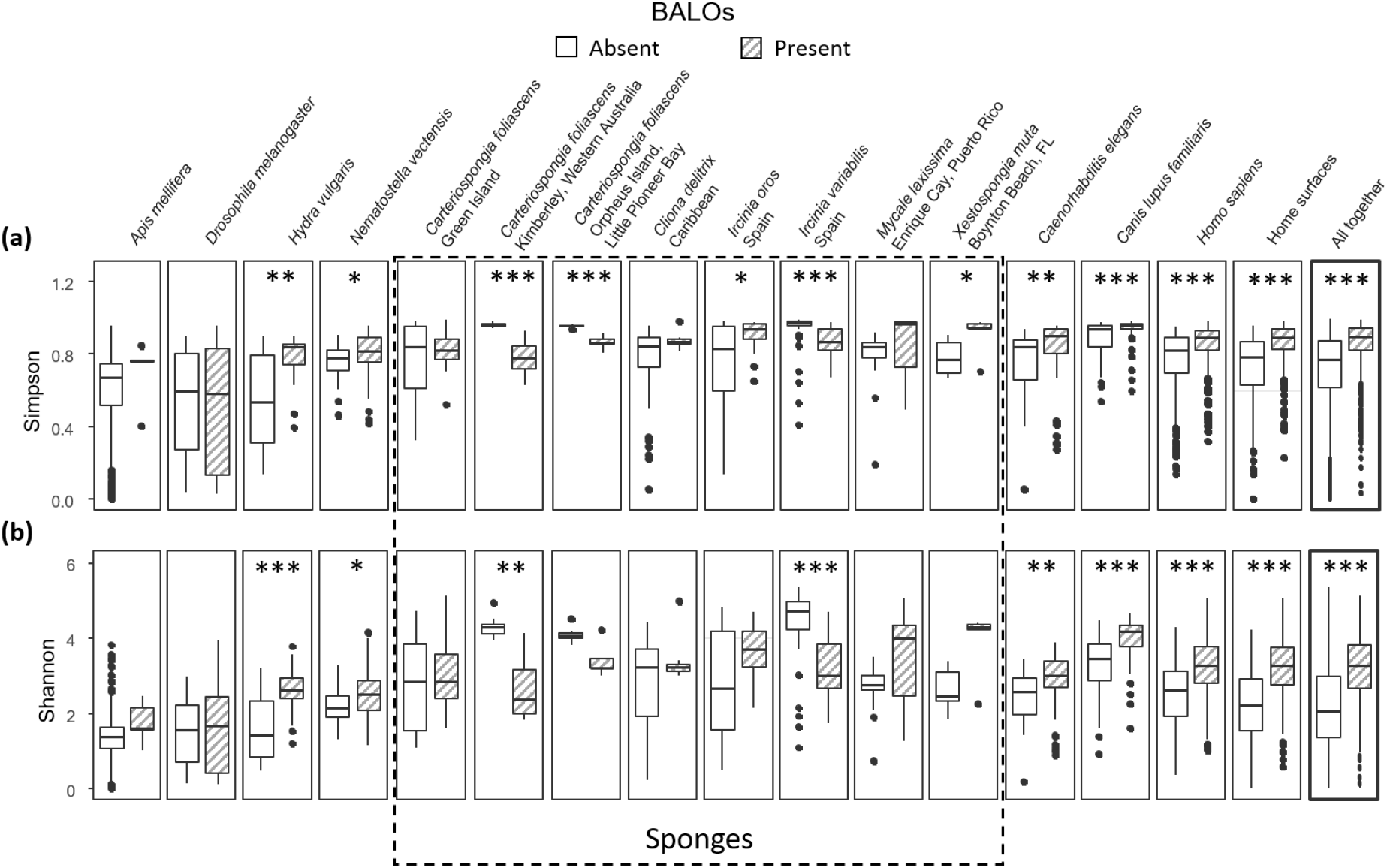
Microbiome alpha-diversity in the presence and absence of BALOs. The Simpson (a) and Shannon (b) diversity is shown for a set of different hosts. Significant differences are indicated by asterisks and were calculated using the Wilcoxon rank sum test. P-values: p<0.001:‘***’, 0.0011>p<0.01:‘**’, 0.011>p<0.05:‘*’. P-values are given in the Table S2.

**Fig. 2.**
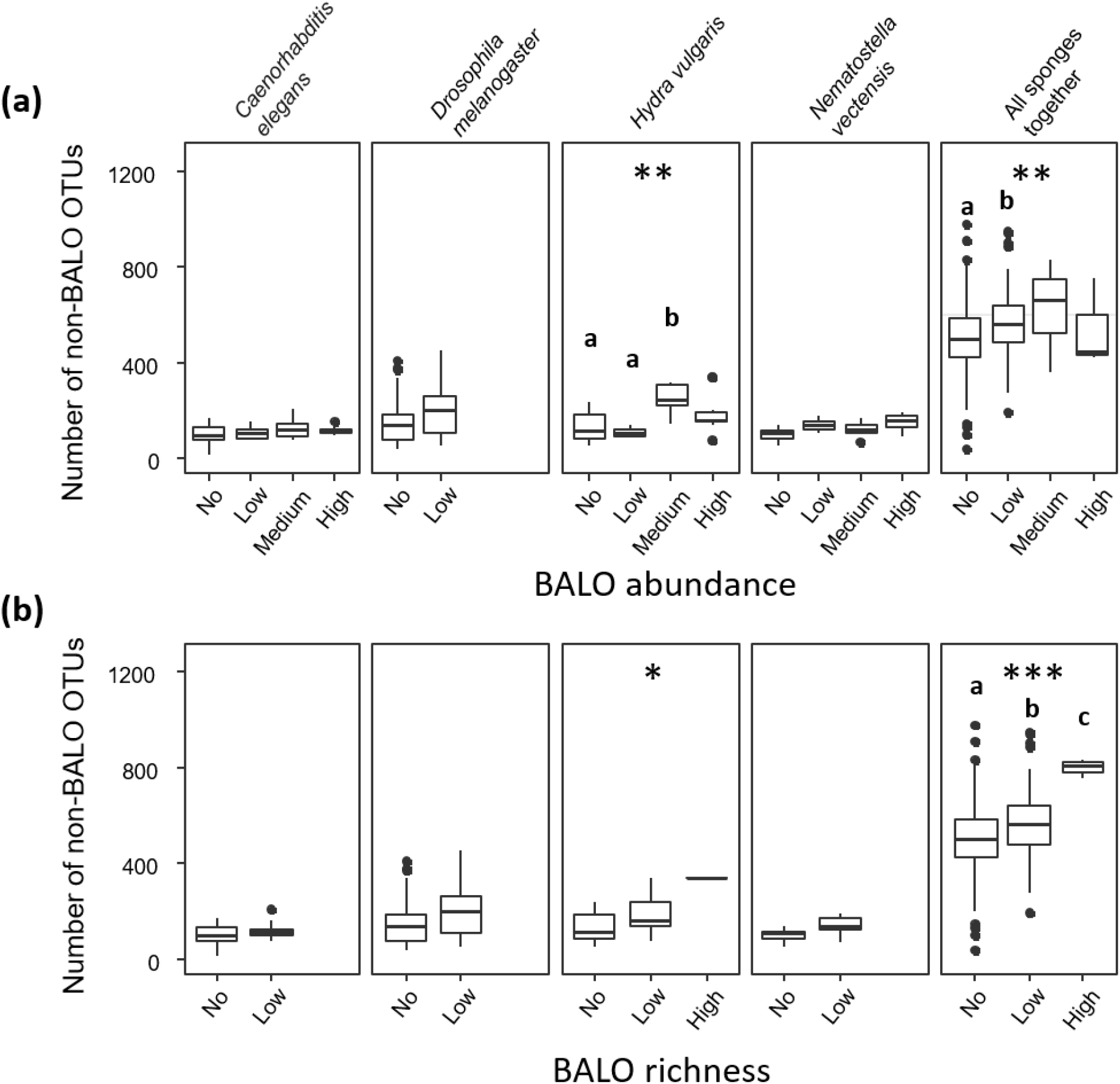
Host microbiome richness measured as number of different non-BALO OTUs with increasing BALO abundance (a) and BALO richness (b). Significant differences are indicated by asterisks and were calculated using the Kruskal-Wallis rank sum test. P-values: p<0.001:‘***’, 0.0011>p<0.01:‘**’, 0.011>p<0.05:‘*’. Significant differences between single categories of BALO abundance and BALO richness are indicated by different letters and were calculated with Dunn’s post hoc test. All P-values are given in the Table S4.

In contrast, variation in microbiome beta-diversity was not linked to the BALOs (Figure 3). At the same time, our PCoA analysis indicated an influence of BALOs on sample clustering for several hosts (especially cnidarians and *C. elegans*). However, the clustering was not independent of sample type, making it impossible to infer the exact cause of clustering from the current data.

**Fig. 3.**
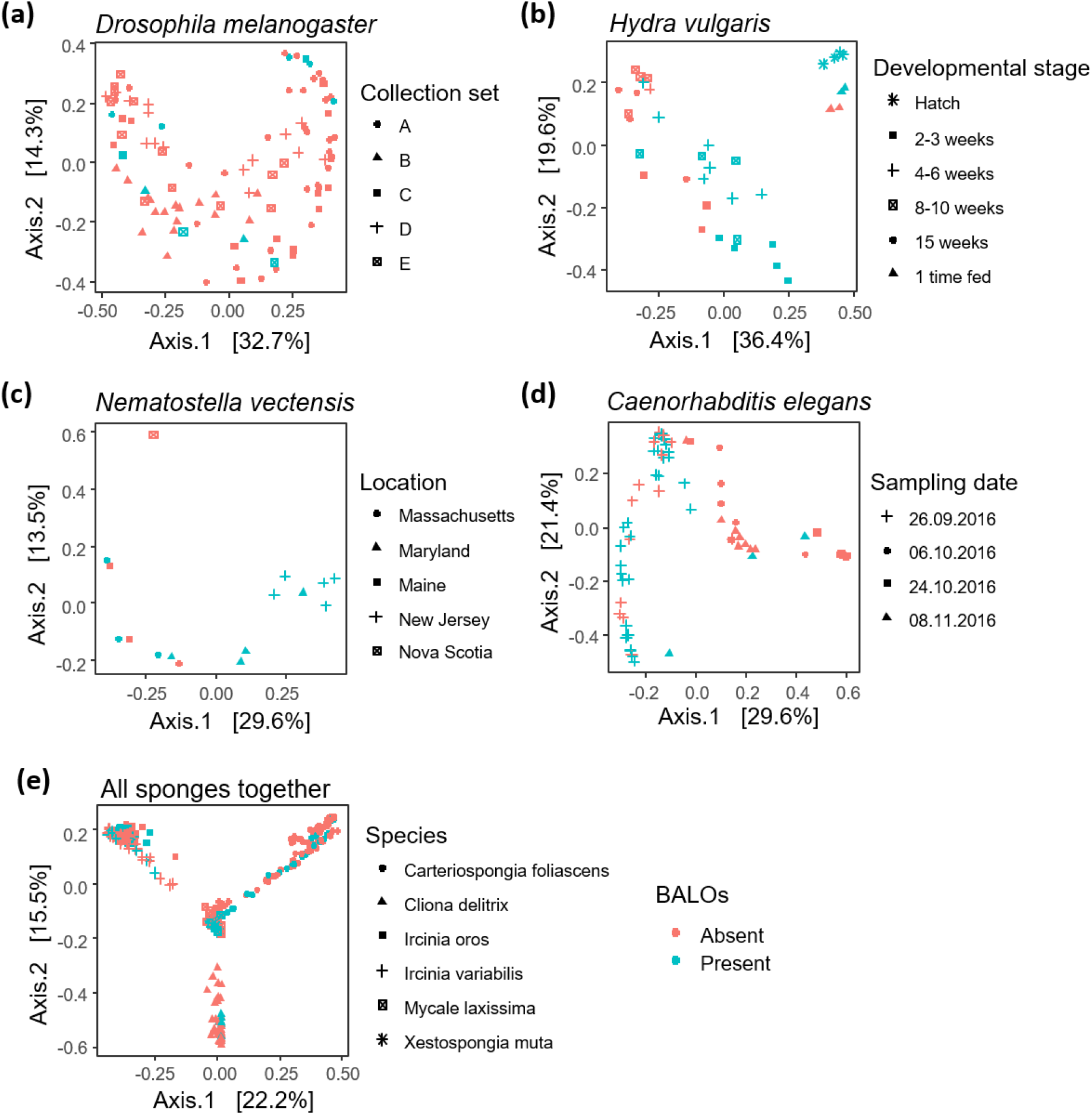
PCoA of microbiome samples from different hosts using Bray Curtis distances. Samples are color-coded by presence and absence of BALOs. Different shapes indicate different sample subsets as indicated by the respective legends.

To exclude that BALO presence is caused by high microbiome diversity as a consequence of sampling effects, we analyzed the complete sponge dataset, additionally including species without BALOs [19]. We found that alpha-diversity *per se* does not predict the presence of BALOs (Table S5 and S6), which is therefore unlikely caused by sampling effects alone.

The loss of top predators has comprehensive effects on community structure [9, 10]. We tested this idea by comparing microbiome alpha-diversities for distinct animal hosts and environments that either lacked or contained a prominent group of microbial predators, the BALOs. With the exception of the considered insects and most sponge species, we found that microbiomes containing BALOs were characterized by a significantly higher alpha-diversity.

In contrast to the overall results, two sponge species showed a negative correlation between BALO presence and microbiome diversity, although not when considering microbiome richness. These results may suggest that BALO-containing sponges harbor a more species-rich, but less even microbiome. Notably, sponges in general possess a comparatively species-rich microbiome (Figure S1). In these cases, evenness may be negatively correlated to richness, consistent with previous observations for plant communities [20] and possibly due to sampling effects, where a superior competitor is more likely present in species-rich communities [1]. A niche-preemption model was previously identified to be the best predictor for the patterns in plant communities [20]. Niche-preemption should favor resource use plasticity among the less competitive species, resulting in lower growth and consequently reduced evenness. In case of the sponges, the negative richness-evenness-relationship might then overshadow the effect of BALOs on microbiome diversity. Temporal effects could additionally explain the higher sponge microbiome diversity in the absence of BALOs. As the sponge data used in this study came from single time point samples, we cannot exclude subsequent changes in the community structure, for example a delayed effect of BALO loss or gain on microbiome diversity. However, the longitudinal data on surface microbiomes [21] indicates that changes in BALO presence/absence are associated with more or less simultaneously occurring changes in OTU richness (Fig. S2).

We found that BALO OTU richness, rather than abundance, is significantly associated with microbiome richness in *H. vulgaris* and the combined set of sponges. Moreover, this significant association between BALO and microbiome richness was only observed when the high BALO richness category could be included. Considering that different BALO strains are known to vary in their range of suitable prey [22], the above results may suggest that a more diverse BALO community is able to prey on a more diverse set of bacteria and thereby reduces the predation pressure on single species, thus increasing microbiome diversity.

Our additional analysis of beta-diversity did not reveal a strong BALO influence on microbiome community structure. Together with the results on alpha-diversity, this may imply that BALO presence is not correlated with a specific community composition and that BALOs survive in a range of differently assembled communities.

Our results from a range of distinct animal hosts and environments point to BALOs as potential drivers of microbiome alpha-diversity, possibly by actively preying on highly abundant species, thereby favoring rare species. Thus, BALOs may be of particular importance for our understanding of the stability and resilience of microbiome ecosystem functions. Our current meta-analysis is, however, based on associations, which can only be indicative of possible causal relationships. An important next step should therefore be a detailed experimental analysis of the exact causal role of BALOs on microbiome diversity and resulting functions. It would be of similar high interest to assess to what extent other kinds of bacterial antagonists, such as phages, or environmental stressors may also influence microbiome diversity and the associated effects. Moreover, it is worth testing whether the interaction between BALOs and other bacteria is additionally shaped by the host immune system, which could cause different dynamics of the BALO-mediated effects within rather than outside host organisms.

Considering that BALOs are not pathogenic to higher organisms [23], have a likely stronger effect on community structure than phages [18], and appear to enhance microbial diversity, they are highly promising candidates for probiotic therapy [24] that aims at restoring disturbed microbiomes and improving host health or ecosystem productivity and stability.

## Acknowledgement

We are grateful for advice from members of the Schulenburg group and the Collaborative Research Center CRC1182 on Origin and Function of Metaorganisms. This work was funded by the CRC1182, projects A4 and B2.

## Supplementary Material

### 1 Supplementary Methods

For our analysis, we randomly selected exemplary host taxa that are representative of distinct taxonomic animal groups, ranging from very simple to more complex hosts and including early branching invertebrates, ecdysozoa, and also vertebrates (Table S1). In addition, we only considered host taxa, for which a single study included at least five samples with and without BALOs - with the exception of the *Nematostella* dataset with only four samples without BALOs. This preselection was performed in order to allow a direct comparison of samples with and without BALOs for each host system or environment. Further, only studies with publicly available OTU tables were selected. Moreover, we considered one study with longitudinal data generated from human mucus, sebum, skin swabs, as well as from different surfaces from their family homes [10]. This data set served to test the stability of the association of BALO presence and bacterial community diversity across time within the same environment.

Several datasets were from microbiomeDB (http://microbiomedb.org/mbio/) and only included relative abundance data, while the remaining data sets also had information on absolute frequencies. The *Caenorhabditis elegans* dataset was produced by us for this study by sampling worms from the Kiel Botanical garden in 2016 at four consecutive time points (one in October, two in September, and one in November). Worm samples were prepared as described previously [1] by using the protocol for "natural worm” microbiome extraction. DNA was sequenced using the Miseq platform and the primers 515f-806r to sequence the V4 region of the 16S rRNA gene. Original sequence data are available from the European Nucleotide Archive (accession number PRJEB30476). Sequence reads were analyzed using Mothur v. 1.39.5 [2] as described in the Miseq SOP (https://www.mothur.org/wiki/MiSeq_SOP) and the SILVA reference database version 128. OTU clustering was based on 97% sequence identity. Samples with BALOs were categorized based on their abundance (i.e., high (11-227 reads), medium (6-10), low (1-5), and no reads) and richness (high (5-7), low (1-4), and no). We compared microbiome alpha-diversity in the presence and absence of BALOs using two-sample Wilcoxon rank sum tests to account for outliers. We assessed the influence of BALO abundance or richness on microbiome richness with the Kruskal-Wallis rank sum test and Dunn’s post hoc test with p-value adjustment using fdr. Beta-diversity was measured using Bray Curtis distance on relative abundance and visualized using PCoA of the 500 most abundant OTUs. Sponge samples were analyzed using Fisher’s exact test and the Wilcoxon rank sum test to test for an association between BALO presence/absence and microbiome alpha-diversity, either as categorical or continues variable. All statistical analyses were performed in R [3] using phyloseq [4] and vegan [5].

### 2 Supplementary Figures and Tables

**Table S1:** Summary of the considered and analyzed studies.

| Study | Host body site | Seq. platform | Environment of host | N samples | Normalization | Further information |
| --- | --- | --- | --- | --- | --- | --- |
| ***Caenorhabditis elegans* (Nematoda)** |  |  |  |  |  |  |
| **This study\*** | Gut | Miseq (V4) | Natural isolates from compost heaps | 73 | 4986 reads per sample | Time series |
| ***Nematostella vectensis* (Cnidaria)** |  |  |  |  |  |  |
| **[6]** | Whole animals | 454 (V2) | Natural isolates, but maintained in the lab for 10 years as clonal lines | 16 | 3000 reads per sample | Microbiome diversity of species sampled along the US east coast |
| **Six sponge species** |  |  |  |  |  |  |
| **[7]** | Random sponge pieces | Hiseq (V4) | Natural isolates from different sites | 315 samples | No normalization | Different species and different sampling sites |
| ***Drosophila melanogaster* (Insecta, Diptera)** |  |  |  |  |  |  |
| **[8]** | Whole flies | Miseq (V3-V4) | Samples taken from various kitchen | 79 | 1200 reads per sample | Only adult flies |
| ***Apis mellifera* (Insecta, Hymenoptera)** |  |  |  |  |  |  |
| **Unpublished, Dominguez-Bello** | whole head, whole larva, whole pupa, whole gut | Unknown | Unknown | 383 | Unknown | From microbiomeDB, different functional guilds and developmental stages, effect of Tetracycline application |
| ***Canis lupus familiaris* (Vertebrata, Mammalia)** |  |  |  |  |  |  |
| **[9]** | Sebum | Illumina GAllx (V2) | Different homes | 145 | 5000 reads per sample | From microbiomeDB, most BALOs in sebum und mucus |
| **Home surfaces** |  |  |  |  |  |  |
| **[10]** | Different surfaces from family homes | Hiseq (V4) | Different homes | 690 | 2500 reads per sample | From microbiomeDB |
| ***Homo sapiens* (Vertebrata, Mammalia)** |  |  |  |  |  |  |
| **[10]** | Mucus, sebum, skin swabs | Hiseq (V4) | Different homes | 910 | 2500 reads per sample | From microbiomeDB |
| ***Hydra vulgaris* (Cnidaria)** |  |  |  |  |  |  |
| **[11]** | Whole animals | 454 (V1-V2) | Lab-kept animals | 36 | No normalization | Developmental data |

**Table S2:**
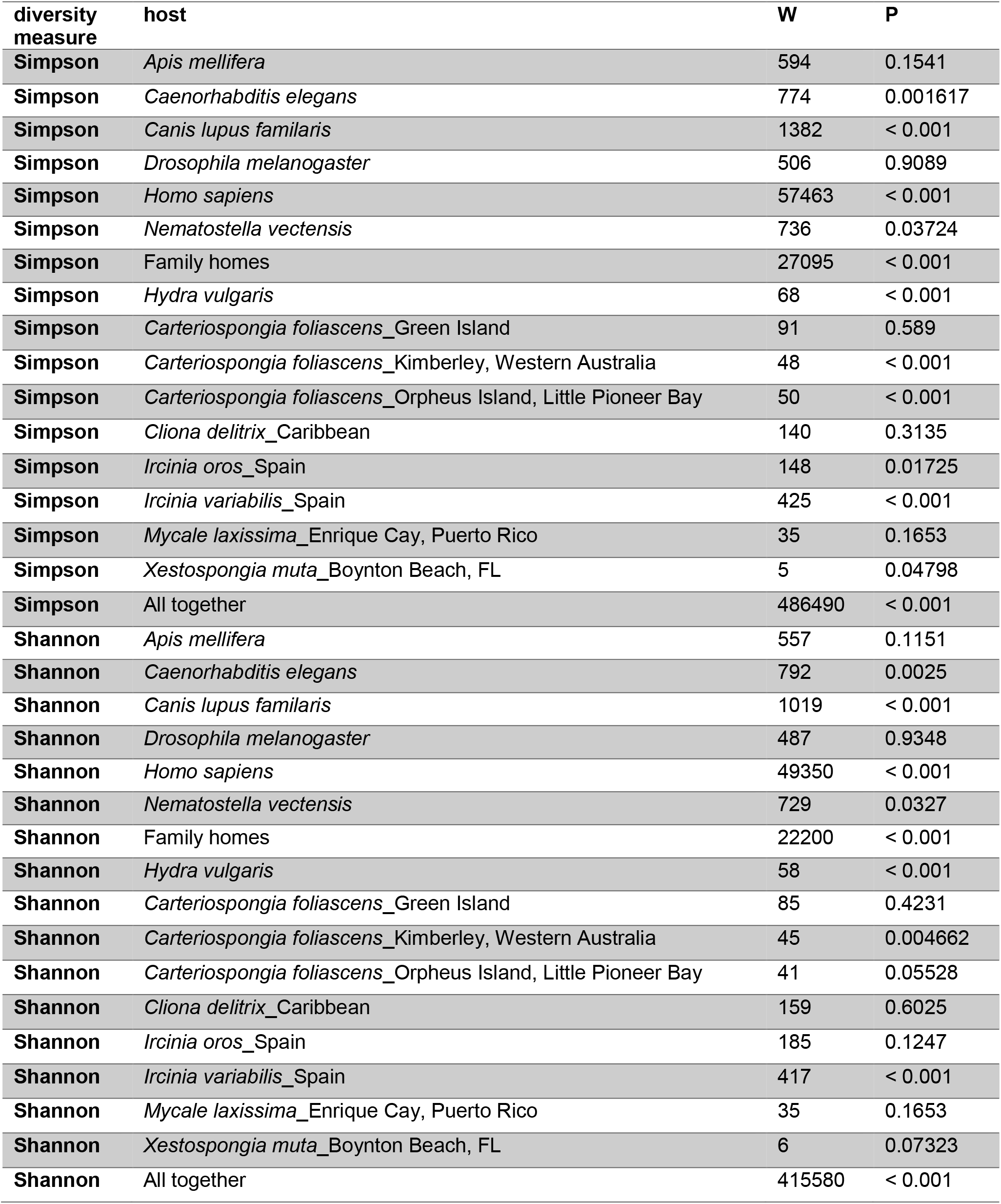
Test statistics of the comparison of microbiome alpha-diversity in the absence and presence of BALOs as shown in Fig. 1.

| diversity<br>measure | host | W | P |
| --- | --- | --- | --- |
| **Simpson** | *Apis mellifera* | 594 | 0.1541 |
| **Simpson** | *Caenorhabditis elegans* | 774 | 0.001617 |
| **Simpson** | *Canis lupus familiaris* | 1382 | < 0.001 |
| **Simpson** | *Drosophila melanogaster* | 506 | 0.9089 |
| **Simpson** | *Homo sapiens* | 57463 | < 0.001 |
| **Simpson** | *Nematostella vectensis* | 736 | 0.03724 |
| **Simpson** | Family homes | 27095 | < 0.001 |
| **Simpson** | *Hydra vulgaris* | 68 | < 0.001 |
| **Simpson** | *Carteriospongia foliascens*_Green Island | 91 | 0.589 |
| **Simpson** | *Carteriospongia foliascens*_Kimberley, Western Australia | 48 | < 0.001 |
| **Simpson** | *Carteriospongia foliascens*_Orpheus Island, Little Pioneer Bay | 50 | < 0.001 |
| **Simpson** | *Cliona delitrix*_Caribbean | 140 | 0.3135 |
| **Simpson** | *Ircinia oros*_Spain | 148 | 0.01725 |
| **Simpson** | *Ircinia variabilis*_Spain | 425 | < 0.001 |
| **Simpson** | *Mycale laxissima*_Enrique Cay, Puerto Rico | 35 | 0.1653 |
| **Simpson** | *Xestospongia muta*_Boynton Beach, FL | 5 | 0.04798 |
| **Simpson** | All together | 486490 | < 0.001 |
| **Shannon** | *Apis mellifera* | 557 | 0.1151 |
| **Shannon** | *Caenorhabditis elegans* | 792 | 0.0025 |
| **Shannon** | *Canis lupus familiaris* | 1019 | < 0.001 |
| **Shannon** | *Drosophila melanogaster* | 487 | 0.9348 |
| **Shannon** | *Homo sapiens* | 49350 | < 0.001 |
| **Shannon** | *Nematostella vectensis* | 729 | 0.0327 |
| **Shannon** | Family homes | 22200 | < 0.001 |
| **Shannon** | *Hydra vulgaris* | 58 | < 0.001 |
| **Shannon** | *Carteriospongia foliascens*_Green Island | 85 | 0.4231 |
| **Shannon** | *Carteriospongia foliascens*_Kimberley, Western Australia | 45 | 0.004662 |
| **Shannon** | *Carteriospongia foliascens*_Orpheus Island, Little Pioneer Bay | 41 | 0.05528 |
| **Shannon** | *Cliona delitrix*_Caribbean | 159 | 0.6025 |
| **Shannon** | *Ircinia oros*_Spain | 185 | 0.1247 |
| **Shannon** | *Ircinia variabilis*_Spain | 417 | < 0.001 |
| **Shannon** | *Mycale laxissima*_Enrique Cay, Puerto Rico | 35 | 0.1653 |
| **Shannon** | *Xestospongia muta*_Boynton Beach, FL | 6 | 0.07323 |
| **Shannon** | All together | 415580 | < 0.001 |

**Table S3:** Test statistics of the comparison of microbiome richness in the presence and absence of BALOs as shown in Fig. S1.

| Host | W | P |
| --- | --- | --- |
| *Caenorhabditis elegans* | 178.5 | 0.06802 |
| *Drosophila melanogaster* | 348 | 0.1102 |
| *Hydra vulgaris* | 88.5 | 0.003041 |
| *Nematostella vectensis* | 11 | 0.1293 |
| All sponges together | 4531.5 | < 0.001 |
| *Carteriospongia foliascens*_Green Island | 144 | < 0.001 |
| *Carteriospongia foliascens*_Kimberley, Western Australia | 70 | 0.9321 |
| *Carteriospongia foliascens*_Orpheus Island, Little Pioneer Bay | 116 | 0.4946 |
| *Cliona delitrix*_Caribbean | 468 | 0.021 |
| *Ircinia oros*_Spain | 518 | < 0.001 |
| *Ircinia variabilis*_Spain | 664 | 0.01112 |
| *Mycale laxissima*_Enrique Cay, Puerto Rico | 220 | 0.9319 |
| *Xestospongia muta*_Boynton Beach, FL' | 42 | 0.1058 |

**Table S4:** Test statistics of the comparison of microbiome richness and BALO abundance and BALO richness as shown in Fig. 2.

| host | category | Kruskal-Wallis<br>Chi-squared | Df | P | Significant Dunn's |
| --- | --- | --- | --- | --- | --- |
| *Caenorhabditis elegans* | BALO abundance | 3.3508 | 1 | 0.06717 |  |
| *Drosophila melanogaster* | BALO abundance | 6.3712 | 3 | 0.09488 |  |
| *Hydra vulgaris* | BALO abundance | 12.795 | 3 | 0.005102 | Medium:Low P = 0.014,<br>No:Medium P = 0.011 |
| *Nematostella vectensis* | BALO abundance | 3.7776 | 3 | 0.2865 |  |
| All sponges together | BALO abundance | 14.573 | 3 | 0.002221 | No:Low P = 0.0033 |
| *Caenorhabditis elegans* | BALO diversity | 4.2926 | 3 | 0.2316 |  |
| *Drosophila melanogaster* | BALO diversity | 2.5684 | 1 | 0.109 |  |
| *Hydra vulgaris* | BALO diversity | 6.7652 | 2 | 0.03396 |  |
| *Nematostella vectensis* | BALO diversity | 2.489 | 1 | 0.1146 |  |
| All sponges together | BALO diversity | 17.87 | 2 | < 0.001 | No:Low P = 0.0032, No:High P =<br>0.0053, Low:High P = 0.0349 |

**Table S5:** Test statistics of the comparison of sponge microbiome alpha-diversity category and BALO presence. Contingency tables are based on the average of the respective value (given in brackets for the different categories) for each species.

|  | Microbiome richness<sup>a</sup> |  | Fisher's Exact Test |  |
| --- | --- | --- | --- | --- |
| | high ($\geq 600$ - 809.25) | low (306.67 - $< 600$) | P | Odds ratio |
| **BALOs present** | 10 | 38 | 0.7356 | 0.79247 |
| **BALOs absent** | 4 | 12 |  |  |

|  | Microbiome Simpson diversity |  | Fisher's Exact Test |  |
| --- | --- | --- | --- | --- |
| | high ($\geq 0.9$) | low (0.38 - $< 0.9$) | P | Odds ratio |
| **BALOs present** | 17 | 31 | 1 | 1.20297 |
| **BALOs absent** | 5 | 11 |  |  |

|  | Microbiome Shannon diversity |  | Fisher's Exact Test |  |
| --- | --- | --- | --- | --- |
| | high (3.5 - 4.74) | low (1.7 - $< 3.5$) | P | Odds ratio |
| **BALOs present** | 23 | 25 | 1 | 1.17976 |
| **BALOs absent** | 7 | 9 |  |  |

<sup>a</sup> Microbiome richness is treated as a categorical variable, being either high or low. Cut-offs for the two groups are indicated.

**Table S6:**
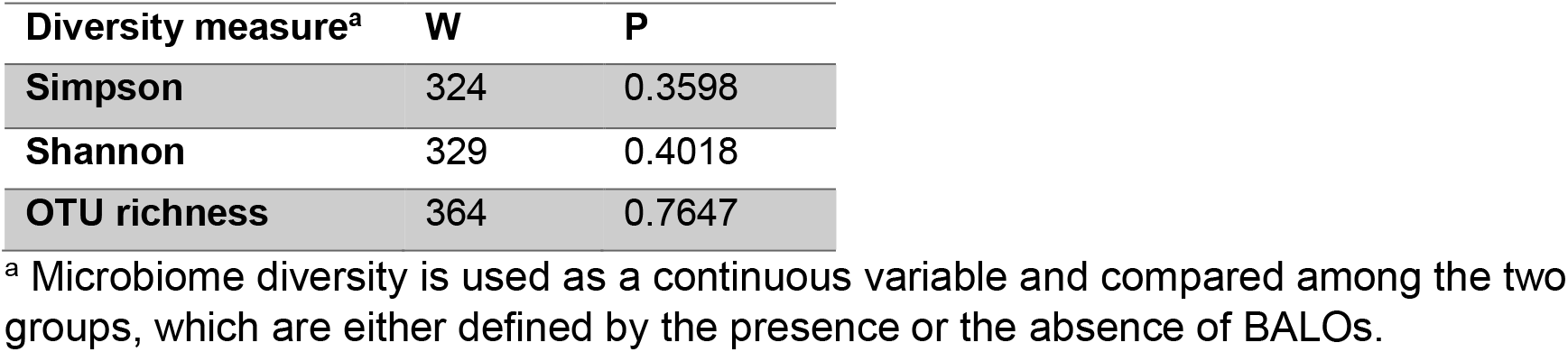
Wilcoxon rank sum test statistics of the comparison of sponge microbiome alpha-diversity in samples either with BALO presence *versus* BALO absence.

**Fig. S1:**
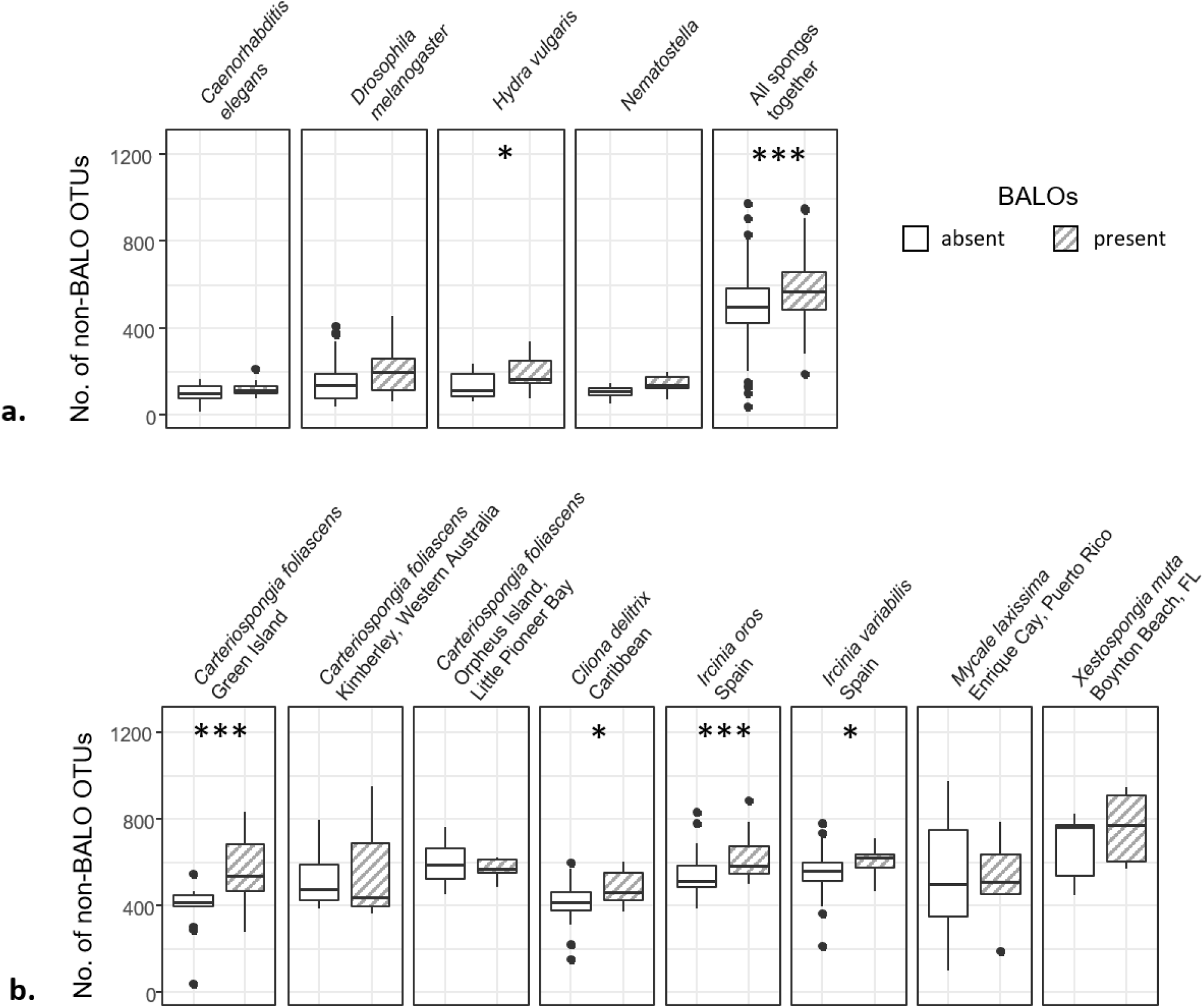
Microbiome richness of different hosts (a) and particular sponge species (b) measured as number of different non-BALO OTUs in the presence and absence of BALOs. Significant differences are indicated by asterisks and were calculated using the Wilcoxon rank sum test. P-values: p<0.001:‘***’, 0.0011>p<0.01:‘**’, 0.011>p<0.05:’*’. P-values are given in the Table S3.

**Fig. S2:**
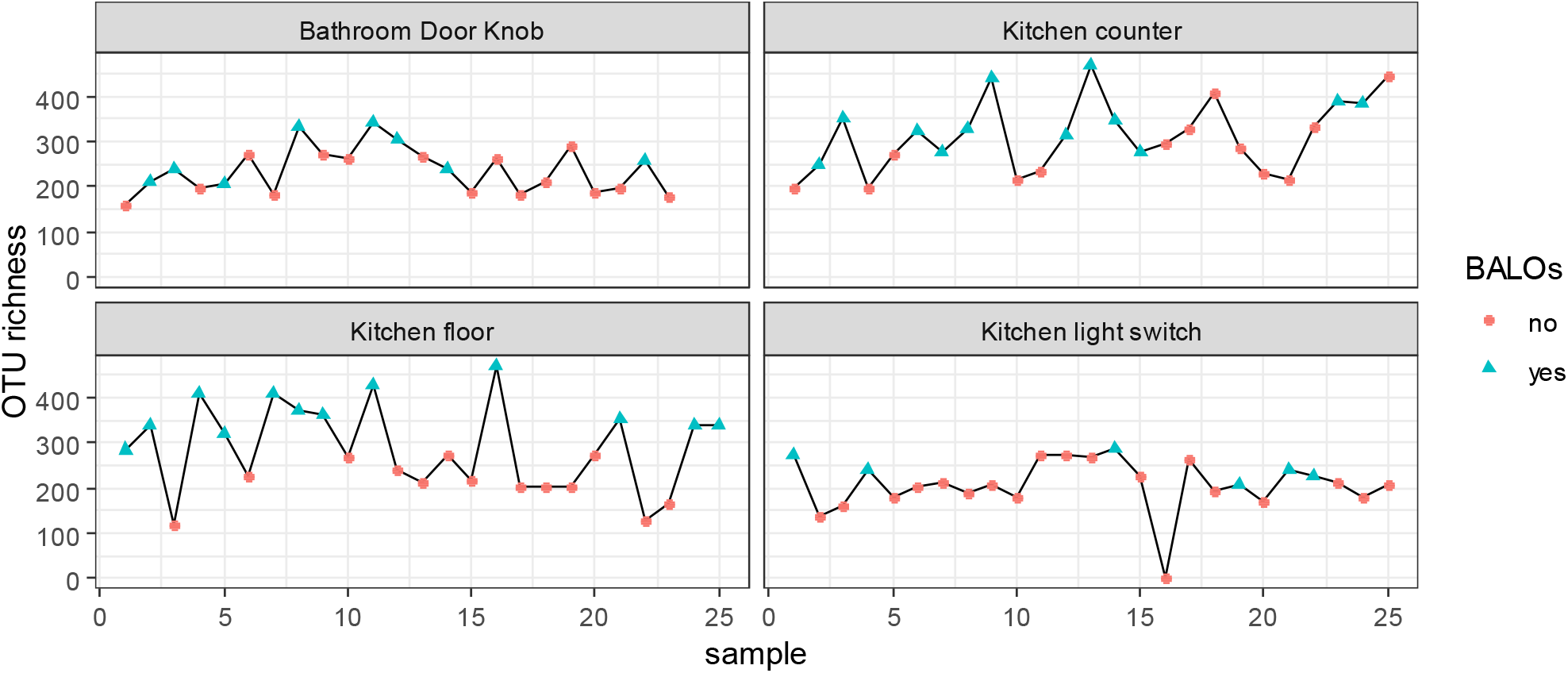
Microbiome richness of longitudinal samples from house 05b from [10]. Samples were taken every other day. Data points are colored and shaped according to the presence or absence of BALOs.

## References

1. Cardinale BJ, Srivastava DS, Duffy JE, et al (2006) Effects of biodiversity on the functioning of trophic groups and ecosystems. Nature 443:989–992. https://doi.org/10.1038/Nature05202

2. Boyer KE, Kertesz JS, Bruno JF Biodiversity effects on productivity and stability of marine macroalgal communities: the role of environmental context. Oikos 118:1062–1072. https://doi.org/10.1111/j.1600-0706.2009.17252.x

3. Power ME, Tilman D, Estes JA, et al (1996) Challenges in the quest for keystones. BioScience 46:609–620. https://doi.org/10.2307/1312990

4. Thompson R, Starzomski BM (2007) What does biodiversity actually do? A review for managers and policy makers. Biodivers Conserv 16:1359–1378. https://doi.org/10.1007/s10531-005-6232-9

5. Naeem S, Li SB (1997) Biodiversity enhances ecosystem reliability. Nature 390:507–509. https://doi.org/10.1038/37348

6. Yachi S, Loreau M (1999) Biodiversity and ecosystem productivity in a fluctuating environment: The insurance hypothesis. Proc Natl Acad Sci 96:1463–1468. https://doi.org/10.1073/pnas.96.4.1463

7. Holling CS (1973) Resilience and stability of ecological systems. Annu Rev Ecol Syst 4:1–23. https://doi.org/10.1146/annurev.es.04.110173.000245

8. Thebault E, Loreau M (2003) Food-web constraints on biodiversity-ecosystem functioning relationships. Proc Natl Acad Sci U S A 100:14949–14954. https://doi.org/10.1073/pnas.2434847100

9. Jabiol J, McKie BG, Bruder A, et al Trophic complexity enhances ecosystem functioning in an aquatic detritus-based model system. J Anim Ecol 82:1042–1051. https://doi.org/10.1111/1365-2656.12079

10. Gessner MO, Swan CM, Dang CK, et al (2010) Diversity meets decomposition. Trends Ecol Evol 25:372–380. https://doi.org/10.1016/j.tree.2010.01.010

11. Leitão RP, Zuanon J, Villéger S, et al (2016) Rare species contribute disproportionately to the functional structure of species assemblages. Proc R Soc B Biol Sci 283:20160084. https://doi.org/10.1098/rspb.2016.0084

12. Callens M, Watanabe H, Kato Y, et al (2018) Microbiota inoculum composition affects holobiont assembly and host growth in Daphnia. Microbiome 6:. https://doi.org/10.1186/s40168-018-0444-1

13. Sockett RE (2009) Predatory lifestyle of Bdellovibrio bacteriovorus. Annu Rev Microbiol 63:523–539. https://doi.org/10.1146/annurev.micro.091208.073346

14. Rotem O, Pasternak Z, Jurkevitch E (2014) Bdellovibrio and like organisms. In: Rosenberg E, DeLong EF, Lory S, et al (eds) The Prokaryotes. Springer Berlin Heidelberg, Berlin, Heidelberg, pp 3–17

15. Iebba V, Santangelo F, Totino V, et al (2013) Higher prevalence and abundance of Bdellovibrio bacteriovorus in the human gut of healthy subjects. Plos One 8:. https://doi.org/10.1371/journal.pone.0061608

16. Negus D, Moore C, Baker M, et al (2017) Predator versus pathogen: How does predatory Bdellovibrio bacteriovorus interface with the challenges of killing Gram-negative pathogens in a host setting? Annu Rev Microbiol 71:441–457. https://doi.org/10.1146/annurev-micro-090816-093618

17. Kandel PP, Pasternak Z, van Rijn J, et al (2014) Abundance, diversity and seasonal dynamics of predatory bacteria in aquaculture zero discharge systems. FEMS Microbiol Ecol 89:149–161. https://doi.org/10.1111/1574-6941.12342

18. Chen H, Laws EA, Martin JL, et al (2018) Relative Contributions of Halobacteriovorax and Bacteriophage to Bacterial Cell Death under Various Environmental Conditions. mBio 9:e01202–18. https://doi.org/10.1128/mBio.01202-18

19. Thomas T, Moitinho-Silva L, Lurgi M, et al (2016) Diversity, structure and convergent evolution of the global sponge microbiome. Nat Commun 7:11870. https://doi.org/10.1038/ncomms11870

20. Zhang H, John R, Peng Z, et al (2012) The relationship between species richness and evenness in plant communities along a successional gradient: A study from sub-alpine meadows of the eastern Qinghai-Tibetan plateau, China. PLoS ONE 7:. https://doi.org/10.1371/journal.pone.0049024

21. Jurkevitch E, Minz D, Ramati B, Barel G (2000) Prey range characterization, ribotyping, and diversity of soil and rhizosphere Bdellovibrio spp. isolated on phytopathogenic bacteria. Appl Env Microbiol 66:2365–71

22. Gupta S, Tang C, Tran M, Kadouri DE (2016) Effect of Predatory Bacteria on Human Cell Lines. PLOS ONE 11:e0161242. https://doi.org/10.1371/journal.pone.0161242

23. Dwidar M, Monnappa AK, Mitchell RJ (2012) The dual probiotic and antibiotic nature of Bdellovibrio bacteriovorus. BMB Rep 45:71–78. https://doi.org/10.5483/BMBRep.2012.45.2.71

24. Mortzfeld BM, Urbanski S, Reitzel AM, et al (2016) Response of bacterial colonization in Nematostella vectensis to development, environment and biogeography. Environ Microbiol 18:1764–1781. https://doi.org/10.1111/1462-2920.12926

25. Adair KL, Wilson M, Bost A, Douglas AE (2018) Microbial community assembly in wild populations of the fruit fly Drosophila melanogaster. ISME J 12:959–972. https://doi.org/10.1038/s41396-017-0020-x

26. Song SJ, Lauber C, Costello EK, et al (2013) Cohabiting family members share microbiota with one another and with their dogs. eLife 2:e00458. https://doi.org/10.7554/eLife.00458

27. Lax S, Smith DP, Hampton-Marcell J, et al (2014) Longitudinal analysis of microbial interaction between humans and the indoor environment. Science 345:1048–1052. https://doi.org/10.1126/science.1254529

28. Franzenburg S, Fraune S, Altrock PM, et al (2013) Bacterial colonization of Hydra hatchlings follows a robust temporal pattern. ISME J 7:781–790. https://doi.org/10.1038/ismej.2012.156

## Supplementary References

1. Dirksen P, Marsh SA, Braker I, et al (2016) The native microbiome of the nematode Caenorhabditis elegans: gateway to a new host-microbiome model. BMC Biol 14:1–16. https://doi.org/10.1186/s12915-016-0258-1

2. Schloss PD, Westcott SL, Ryabin T, et al (2009) Introducing mothur: open-source, platform-independent, community-supported software for describing and comparing microbial communities. Appl Environ Microbiol 75:7537–7541. https://doi.org/10.1128/Aem.01541-09

3. R Core Team (2016) R: A language and environment for statistical computing. R Foundation for Statistical Computing, Vienna, Austria

4. McMurdie PJ, Holmes S (2013) phyloseq: An R package for reproducible interactive analysis and graphics of microbiome census data. PLOS ONE 8:e61217. https://doi.org/10.1371/journal.pone.0061217

5. Dixon P (2003) VEGAN, a package of R functions for community ecology. J Veg Sci 14:927–930. https://doi.org/10.1111/j.1654-1103.2003.tb02228.x

6. Mortzfeld BM, Urbanski S, Reitzel AM, et al (2016) Response of bacterial colonization in Nematostella vectensis to development, environment and biogeography. Environ Microbiol 18:1764–1781. https://doi.org/10.1111/1462-2920.12926

7. Thomas T, Moitinho-Silva L, Lurgi M, et al (2016) Diversity, structure and convergent evolution of the global sponge microbiome. Nat Commun 7:11870. https://doi.org/10.1038/ncomms11870

8. Adair KL, Wilson M, Bost A, Douglas AE (2018) Microbial community assembly in wild populations of the fruit fly Drosophila melanogaster. ISME J 12:959–972. https://doi.org/10.1038/s41396-017-0020-x

9. Song SJ, Lauber C, Costello EK, et al (2013) Cohabiting family members share microbiota with one another and with their dogs. eLife 2:e00458. https://doi.org/10.7554/eLife.00458

10. Lax S, Smith DP, Hampton-Marcell J, et al (2014) Longitudinal analysis of microbial interaction between humans and the indoor environment. Science 345:1048–1052. https://doi.org/10.1126/science.1254529

11. Franzenburg S, Fraune S, Altrock PM, et al (2013) Bacterial colonization of Hydra hatchlings follows a robust temporal pattern. ISME J 7:781–790. https://doi.org/10.1038/ismej.2012.156

